# Potentiating the cross-reactive IFN-γ T cell and polyfunctional T cell responses by heterologous GX-19N DNA booster in mice primed with either a COVID-19 mRNA vaccine or inactivated vaccine

**DOI:** 10.1101/2022.05.29.493923

**Authors:** Yong Bok Seo, Ara Ko, Duckhyang Shin, Junyoung Kim, You Suk Suh, Juyoung Na, Ji In Ryu, Young Chul Sung

## Abstract

Waning vaccine-induced immunity, coupled with the emergence of SARS-CoV-2 variants, has inspired the widespread implementation of COVID-19 booster vaccinations. Here, we evaluated the potentials of the GX-19N DNA vaccine as a heterologous booster to enhance the protective immune response to SARS-CoV-2 in mice primed with either an inactivated virus particle (VP) or mRNA vaccine. We found that in the VP-primed condition, GX-19N enhanced the response of both vaccine-specific antibodies and cross-reactive T-cells to the SARS-CoV-2 variant of concern (VOC) compared to the homologous VP vaccine prime-boost. Under the mRNA-primed condition, GX-19N induced higher vaccine-induced T-cell responses but lower antibody responses than the homologous mRNA vaccine prime-boost. Furthermore, heterologous GX-19N boost induced higher S-specific polyfunctional CD4^+^ and CD8^+^ T cell responses than the homologous VP or mRNA prime-boost vaccinations. Our results provide new insights into booster vaccination strategies for the management of novel COVID-19 variants.

## Introduction

Unprecedented rates of vaccine development occurred in response to the COVID-19 pandemic. As a result, 20 different types of COVID-19 vaccines have been approved for use, with 65.1% and 58.9% of the world’s population receiving at least one dose and full vaccination, respectively^1^. Approximately 4.6 and 3.2 billion doses of virus particle (VP) and mRNA vaccines were delivered globally, respectively^2^.

Considering the increasing prevalence of COVID-19 and the duration of vaccine efficacy, many countries are implementing COVID-19 vaccine-booster programs. Although boosters increased the vaccine-induced immune response (i.e., neutralizing antibody response), the neutralizing antibody titers against the emerging variants were significantly lower than those against the SARS-CoV-2 wild-type^3 4 5 6^. Because the initial boosters were based on the SARS-CoV-2 wild-type sequence, their efficacy inducing a potent neutralizing antibody response to mutated sequences is limited. Therefore, an additional booster shot is currently required approximately 6 months after the completion of a vaccination. To overcome the limitations of SARS-CoV-2 wild-type-based vaccine booster shots, variant-specific vaccines are continuously being developed^7 5^. Nevertheless, the rationale for the SARS-CoV-2 wild-type-based COVID-19 vaccine booster strategy is to prevent hospitalization due to infection rather than preventing infection itself, and the T-cell response rather than the neutralizing antibody response is considered to be the main factor preventing hospitalization. Unlike neutralizing antibody responses, T cell recognition appears to be broadly cross-reactive against variants of concern (VOCs)^8^.

Several vaccine platforms have been developed for COVID-19 vaccines, and clinical results have shown that different types of immune responses are induced by each. The COVID-19 protein subunit vaccine induces an antibody-rather than a T cell-associated immune response^9 10^; the COVID-19 VP vaccine induces and antibody-associated immune response^11 12^; and the COVID-19 viral vector vaccine induces both antibody-and T cell-associated immune responses^13 14^. Similarly, COVID-19 mRNA vaccines effectively induce antibody and T cell responses^15 16^. GX-19N, which is being developed as a COVID-19 DNA vaccine, effectively induces a T cell response with a marginal antibody response^17^.

In this study, we evaluated the performance of GX-19N DNA vaccine booster-shot regimens in mice primed with either the COVID-19 mRNA or VP vaccine. We demonstrated for the first time that the heterologous GX-19N DNA-boosting vaccination induced a much stronger T cell response than either homologous mRNA or VP prime-boost vaccinations. Furthermore, we demonstrated that polyfunctional CD4+ and CD8+ T cell responses were also increased by the heterologous GX-19N DNA-boosting vaccination.

## Result

### Heterologous GX-19N DNA boosting vaccination induced a higher and lower antibody response than homologous VP and mRNA prime-boost, respectively

To investigate the potential of heterologous GX-19N DNA booster vaccination, SARS-CoV-2 mRNA-or VP-primed BALB/c mice were vaccinated with either a homologous-or heterologous-primed GX-19N DNA booster (Figure 1A). The VP vaccine primed-GX-19N induced a much higher S_RBD_-specific antibody response than the homologous VP vaccine prime-boost regimen did, whereas the homologous mRNA vaccine prime-boost regimen induced a significantly higher S_RBD_-specific antibody response than the mRNA vaccine primed-GX-19N did (Figure 1B, C). The ratio of IgG2a to IgG1 antibody titer tended to increase after the GX-19N vaccination, indicating an enhancement of Th1-polarized immunity, consistent with previous reports (Figure 1D, E)^18 19^.

**Figure 1.**
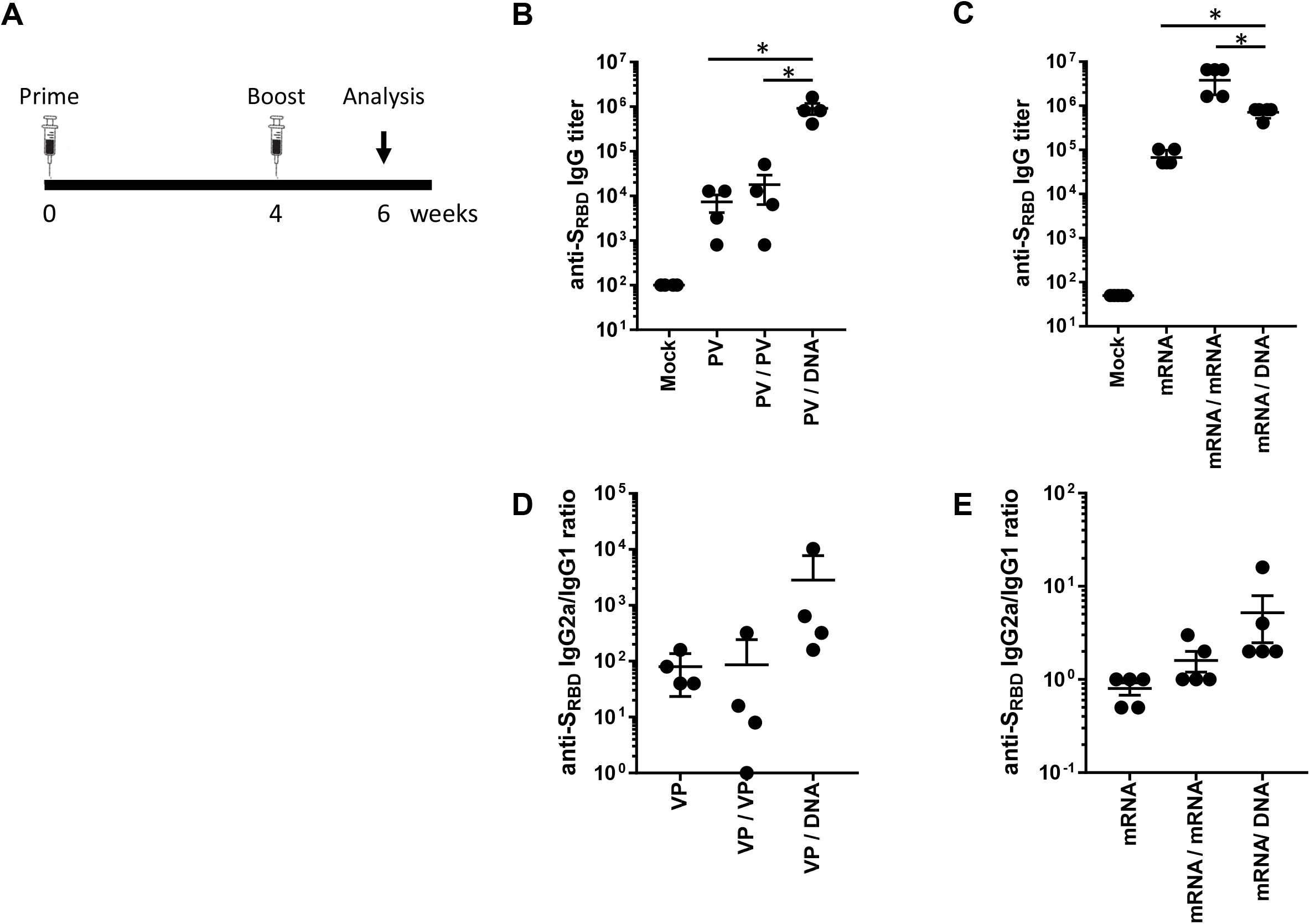
Humoral response to SARS-CoV-2 S_RBD_ antigen after homologous and heterologous prime-boost vaccination. (A) BALB/c mice (n = 4–5/group) were immunized at weeks 0 and 4; serum antibody responses were measured 2 weeks after the last immunization. Graphs show (B, C) SARS-CoV-2 S_RBD_-specific IgG titers and (D, E) ratios of S_RBD_-specific IgG2a to IgG1 titers. Individual mice are represented by a single data point. *P*-values were determined using a two-tailed Student’s *t*-test. *p < 0.05.

In addition, we evaluated neutralizing antibody responses to Wuhan and VOCs (B.1.351 and B.1.617.2) using the surrogate virus neutralization test (sVNT), which is highly correlated with the conventional VNT (cVNT) and pseudovirus-based VNT (pVNT)^20^. In the case of VP vaccine priming, the GX-19N induced a significantly higher neutralizing antibody titer than the homologous VP-boosting vaccination. The neutralizing antibody titer obtained through the heterogeneous regimen was 76.1-fold higher for the Wuhan (mean = 14 vs. 1,076 sVNT_20_ titer), 53.8-fold higher for B.1.351 (mean = 10 vs. 538 sVNT_20_ titer), and 76.1-fold higher for B.1.617.2 (mean = 14 vs. 1,076 sVNT_20_ titer) variants than that obtained with the homologous prime-boost regimen (Figure 2A, B, C). In the case of mRNA vaccine priming, the homologous mRNA-boosting vaccination induced significantly higher neutralizing antibody titers than the GX-19N vaccination. The neutralizing antibody titer obtained through the homologous prime-boost regimen was 2.2-fold higher for the Wuhan (mean = 1,280 vs. 576 sVNT_20_ titer), 2.5-fold higher for B.1.351 (mean = 1,280 vs. 512 sVNT_20_ titer), and 3.2-fold higher for B.1.617.2 (mean = 2,688 vs. 848 sVNT_20_ titer) variants than that obtained with the heterologous prime-boost regimen (Figure 2D, E, F).

**Figure 2.**
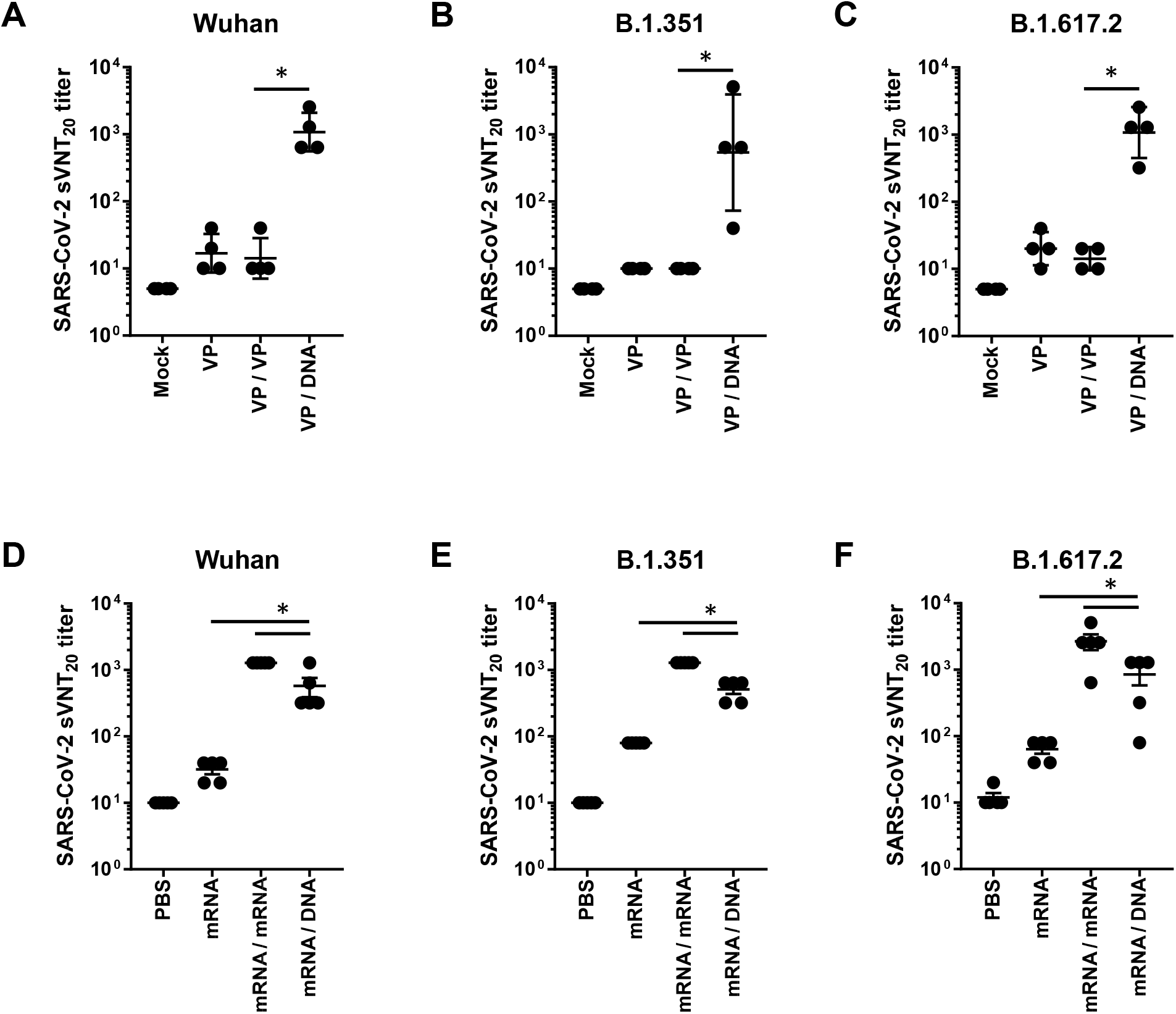
Neutralizing antibody responses against SARS-CoV-2 variants after homologous and heterologous prime-boost vaccination. BALB/c mice (n = 4–5/group) were immunized at weeks 0 and 4; serum antibody responses were measured 2 weeks after the last immunization. Sera from vaccinated mice were tested for (A, D) sVNT^20^ titers in Wuhan, (B, E) B.1.351, (C, F) and B.1.617.2. Individual mice are represented by a single data point. *P*-values were determined using a two-tailed Student’s *t*-test. ns, not significant; *p < 0.05.

### Heterologous GX-19N DNA boosting vaccination induces a higher cross-reactive T cell response against SARS-CoV-2 VOCs than the homologous VP or mRNA prime-boost regimen

T cell responses were evaluated after homologous-or heterologous-primed GX-19N DNA boosting regimens in SARS-CoV-2 mRNA-or VP-primed BALB/c mice. Compared with the homologous VP prime-boost regimen, the GX-19N vaccination increased the T cell response by 21.1-fold (Figure 3A). As expected, we observed similar levels of cellular responses to B.1.351 (mean = 1,644 SFUs/10^6^ splenocytes) and B.1.617.2 (mean = 1,753 SFUs/10^6^ splenocytes) spike peptides (Figure 3B). This is consistent with previous findings of cellular immunity being relatively unimpaired by VOCs compared with neutralizing antibody responses^21^. Similar to the results for VP vaccine-primed mice, the GX-19N increased T cell responses by 2.3-fold compared to the homologous mRNA prime-boost vaccination (Figure 3C).

**Figure 3.**
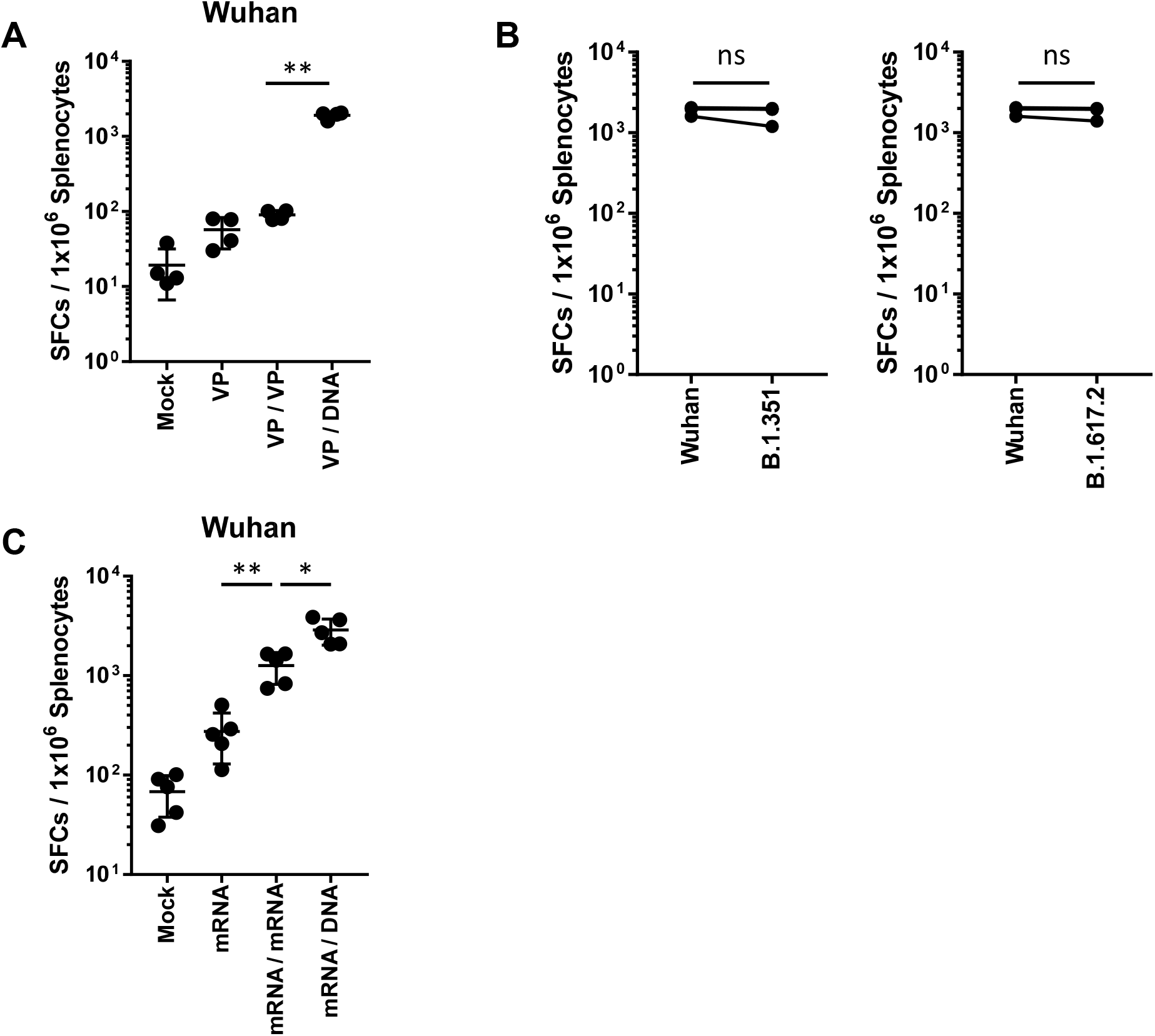
SARS-CoV-2 spike-specific T cell response against SARS-CoV-2 variants after homologous and heterologous prime-boost vaccination. BALB/c mice (n = 4–5/group) were immunized at weeks 0 and 4; T-cell response was measured by IFN-γ ELIspot in splenocytes stimulated with peptide pools spanning the SARS-CoV-2 spike protein. Spike-specific T cell responses were measured in (A, C) Wuhan, (B, D) B.1.351, and B.1.6172. Individual mice are represented by a single data point. *P*-values were determined using a two-tailed Student’s *t*-test. ns, not significant; **p < 0.01.

### Heterologous GX-19N DNA boosting vaccination induces a superior polyfunctional T cell response than the homologous mRNA prime-boost

The quality of the S-specific T cell responses was characterized by analyzing the pattern of cytokine production (i.e., IFN-γ, TNF-α, and/or IL-2) (Figure 4A, Figure 5A). In the S-specific CD4^+^ T cell response, higher IFN-γ^+^TNF-α^+^ or IFN-γ^+^TNF-α^+^IL-2^+^ polyfunctional T cells were induced by the GX-19N than by homologous mRNA prime-boost regimens. The proportion of polyfunctional T cells among cytokine-producing cells was also increased by the GX-19N regimen (mRNA/mRNA, 65% vs. mRNA/GX-19N DNA, 87%) (Figure 4B). An S-specific multifunctional CD8^+^ T cell response was also generated. Similar to CD4^+^ T cell responses, the proportion of multifunctional CD8^+^ T cells among cytokine-producing cells was increased by the GX-19N regimen (mRNA/mRNA, 48% vs. mRNA/GX-19N DNA, 80%) (Figure 5B).

**Figure 4.**
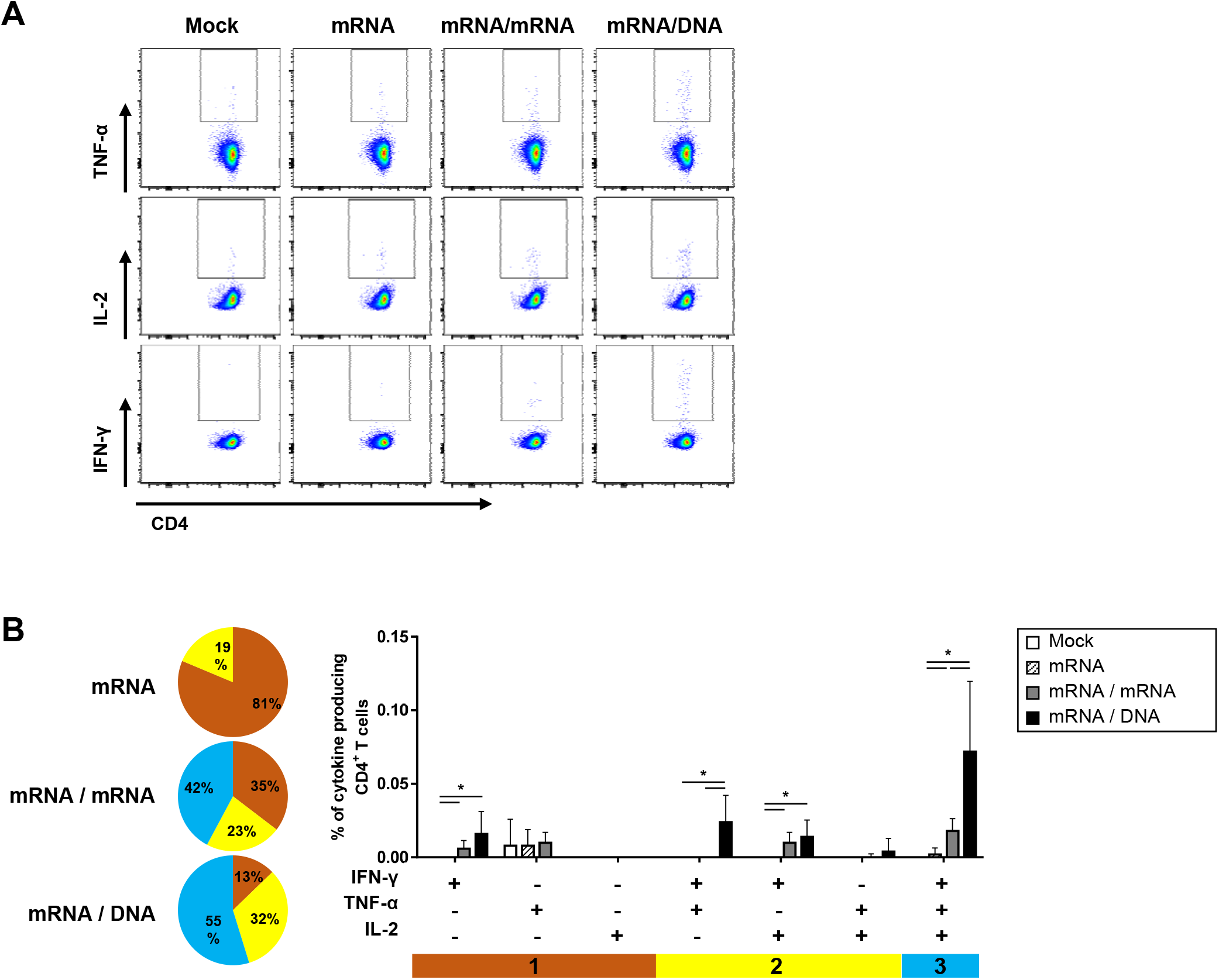
Polyfunctionality of vaccine-induced CD4^+^ T cell response after homologous and heterologous prime-boost vaccination. BALB/c mice (n = 4/group) were immunized at weeks 0 and 4; CD4^+^ T-cell response was measured by ICS in splenocytes stimulated with peptide pools spanning the SARS-CoV-2 spike protein. (A) Vaccine-induced CD4^+^ T cell response. (B) Polyfunctionality of vaccine-induced CD4^+^ T cell responses based on every possible combination of functions. Pie graph sections represent the fraction of T-cells positive for a given number of functions. *P*-values were determined using a two-tailed Student’s *t*-test. ns, not significant; **p < 0.01.

**Figure 5.**
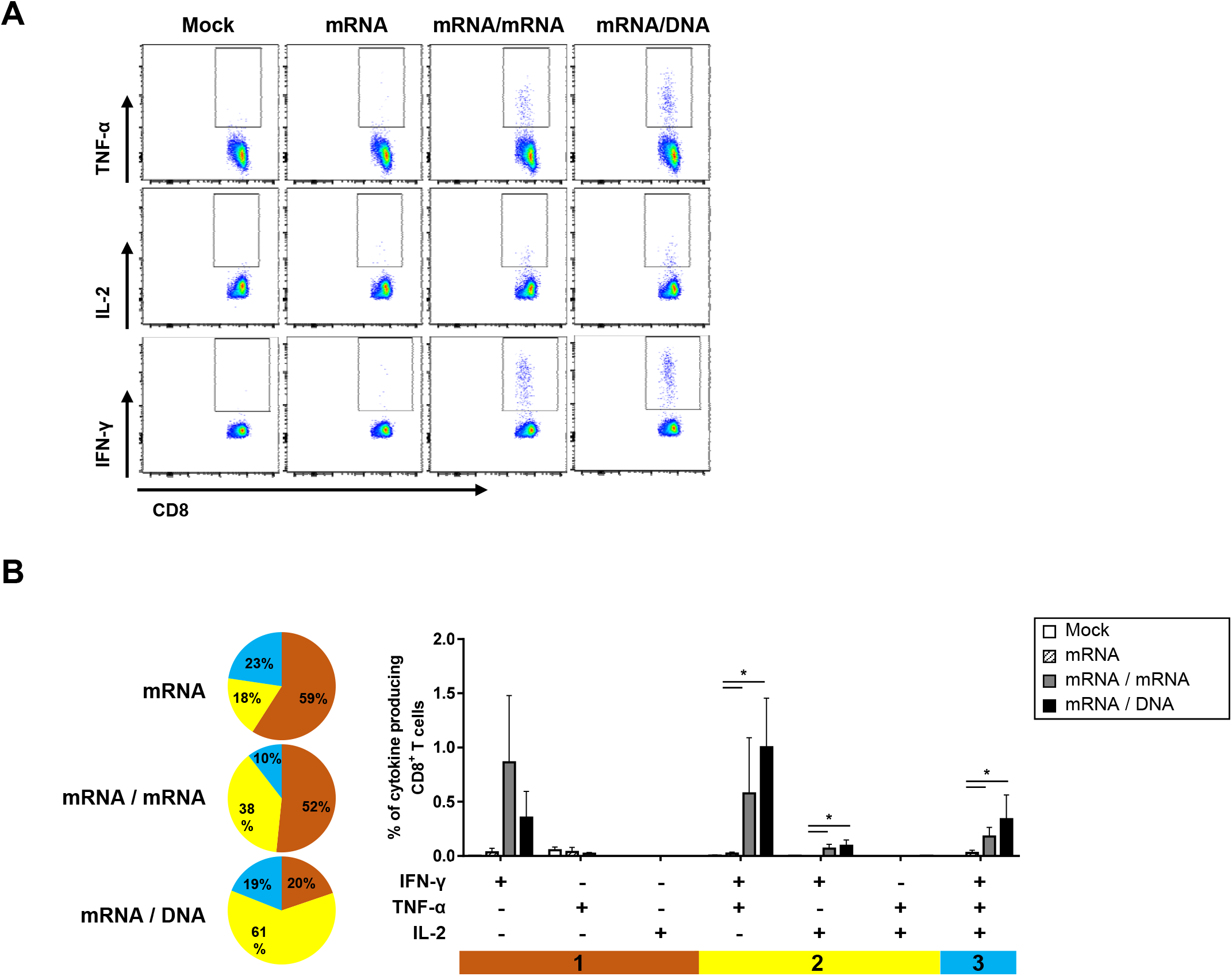
Polyfunctionality of vaccine-induced CD8^+^ T cell response after homologous and heterologous prime-boost vaccination. BALB/c mice (n = 4/group) were immunized at weeks 0 and 4; CD8^+^ T-cell response was measured by ICS in splenocytes stimulated with peptide pools spanning the SARS-CoV-2 spike protein. (A) Vaccine-induced CD8^+^ T cell response. (B) Polyfunctionality of vaccine-induced CD8^+^ T cell responses based on every possible combination of functions. Pie graph sections represent the fraction of T-cells positive for a given number of functions. *P*-values were determined using a two-tailed Student’s *t*-test. ns, not significant; **p < 0.01.

## Discussion

The rapid decline of vaccine-induced immunity against SARS-CoV-2, especially that of humoral immunity against its variants, has led to an increase in infection rates, which prompted the development of vaccine booster doses. As a result of many clinical studies on homologous or heterologous booster doses using current COVID-19 vaccines, the use of booster shots for COVID-19 mRNA has been approved^22 3 6^. The COVID-19 booster vaccine has the advantage of providing substantially increased protection against severe infections and has reduced the rates of hospitalization and death globally^23^. However, the efficacy of the COVID-19 booster dose against SARS-CoV-2 infection remains low (30% or less)^4^. T cells are expected to be particularly important in enhancing protection against severe COVID-19 infections after booster doses. Mounting evidence suggests that T-cell contributions to the host immune response are required for early, broad, and durable protection from SARS-CoV-2, especially with regards of new VOCs^24,25 26 27 28 29^. However, taking two or more booster shots with COVID-19 mRNA vaccines every few months may pose potential risks, including anaphylaxis. Reactions with anaphylactic features following administration of both COVID-19 mRNA vaccines have been reported in the United Kingdom, United States, Japan, and elsewhere^30 31 32 33^. The incidence of these reactions may be higher with these than with other vaccines, such as protein subunit or VP vaccines. Polyethylene glycol (PEG), one of the lipid components of the COVID-19 mRNA vaccine, is hypothesized to be the cause of IgE-mediated anaphylactic reactions to medications, bowel preps, or laxatives containing PEG^34 35^. In addition, it is presumed that the cause of anaphylaxis is a complement activation-related pseudoallergy (CARPA), in which the pre-existing IgG or IgM antibody to PEG activates complementarily, generating anaphylatoxins (C3a, C4a, and C5a), and causing mast cell degranulation^36^. There is also an increased risk of myocarditis. Based on reports of passive surveillance in the United States, the risk of myocarditis after vaccination with COVID-19 mRNA increased across multiple age and sex strata, and was more prevalent after the second dose of the vaccination^37^. This suggests that repeated booster doses of the COVID-19 mRNA vaccine increase the risk of developing myocarditis. Finally, there is an increased likelihood of antibody-dependent enhancement (ADE). Although no serious ADE concerns have arisen with the COVID-19 vaccine, studies on COVID-19 patients have reported a high potential for ADE ^38 39^. Repeated booster shots of the SARS-CoV-2 wild-type sequence-based COVID-19 mRNA vaccine, which can induce strong humoral and T cell responses, can increase non-neutralizing antibody responses to SARS-CoV-2 variants. Given that ADE are mediated by non-neutralizing antibody responses, the likelihood of developing ADEs may gradually increase. Taken together, these findings suggest that an ideal COVID-19 vaccine booster should be safe and T cell-oriented. In this study, we investigated the performance of the GX-19N DNA vaccine, which demonstrated an excellent safety profile and T cell-biased immune response in a previous clinical study^17^, as a booster for mRNA or VP vaccines.

Interestingly, the heterologous GX-19N DNA booster vaccination significantly increased the vaccine-induced T cell response, more so than the homologous mRNA booster. In contrast, the induction of antibody response was lower in the GX-19N than in the homologous mRNA booster regimen. This may be due to the different responses between Th1 (type 1 T helper) and Th2 among the different vaccines used in this study. It is known that the Th1 response induces a cell-mediated response, whereas the Th2 response is related to the humoral immune response^40^. COVID-19 mRNA vaccines can induce an immune response that is unbiased for Th1 or Th2 responses in mice^41 42^. In contrast, the GX-19N DNA vaccine induced a Th1-predominant response in mice^43^. Considering that Th1 and Th2 responses are distinctly related to cellular and humoral immune responses, the immune response induced by the mRNA vaccine prime vaccination would be biased toward the Th1 response induced by the GX-19N boost vaccination. As a result, the heterologous GX-19N DNA booster induced a high T cell response but a low antibody response compared to the mRNA booster vaccination. The higher ratio of IgG2a to IgG1 antibody titers observed using the GX-19N booster compared to the homologous mRNA prime-boost support the above explanation.

## Methods

### Vaccines

The COVID-19 GX-19N DNA vaccine, consisting of GX-19 and GX-21 at a ratio of 1:2, was constructed by inserting the antigen genes of SARS-CoV-2 into a pGX27 vector. GX-19 (pGX27-S_ΔTM_/IC) contains the SARS-CoV-2 spike (S) gene lacking the transmembrane (TM)/intracellular (IC) domain, and GX-21 (pGX27-S_RBD_-F/NP) is designed to express the fusion protein of the receptor-binding domain (RBD) of the spike protein, the T4 fibritin C-terminal foldon (S_RBD_-Foldon), and the nucleocapsid protein (N). S, S_RBD_-Foldon, and N are preceded by the secretory signal sequence of tissue plasminogen activation (tPA). The inactivated SARS-CoV-2 vaccine produced from Vero cells contained 4 μg of viral antigens and 0.225 mg of aluminum hydroxide adjuvant in a 0.5-mL dose. The mRNA vaccine was encapsulated in a lipid nanoparticle through a modified ethanol-drop nanoprecipitation process. Briefly, ionizable, structural, helper, and PEG lipids were mixed with mRNA in acetate buffer at a ratio of 2.5:1 (lipids:mRNA). The mixture was neutralized with Tris-Cl, sucrose was added as a cryoprotectant, and the final solution was sterile filtered and stored frozen at –70 °C until further use.

### Mouse immunizations

Female BALB/c mice aged 6–8 weeks (Central Lab Animal) were intramuscularly immunized with 0.4 μg/animal VP vaccine (total volume of 50 μL, adjusted with PBS) at week 0. At week 4, the same mice were injected with 12 μg/animal GX-19N booster (total volume of 50 μL, adjusted with PBS) into the tibialis anterior muscle with *in vivo* electroporation using an OrbiJector® system (SL VaxiGen Inc.). Mice were sacrificed two weeks after the final immunization.

### Antigen binding ELISA

The serum collected at each time point was evaluated for binding titers. In this assay, 96-well ELISA plates (NUNC) were coated with 1 μg/mL recombinant SARS-CoV-2 spike RBD-His protein (Sino Biological 40592-V08B) in PBS overnight at 4 °C. The plates were washed three times with 0.05% PBST (Tween 20 in PBS) and blocked with 5% skim milk in 0.05% PBST (SM) for 2–3 h at room temperature. The sera were serially diluted in 5% SM, added to the wells, and incubated for 2 h at 37 °C. Following incubation, the plates were washed five times with 0.05% PBST and then incubated with horseradish peroxidase (HRP)-conjugated anti-mouse IgG (Jackson ImmunoResearch Laboratories 115-035-003), IgG1 (Jackson ImmunoResearch Laboratories 115-035-205), or IgG2a (Jackson ImmunoResearch Laboratories 115-035-206) for 1 h at 37 °C. After the final wash, the plates were developed using TMB solution (Surmodics TMBW-1000-01), and the reaction was stopped with 2N H_2_SO_4_. The plates were analyzed at 450 nm using a SpectraMax Plus384 (Molecular Devices).

### Surrogate virus-neutralization assay

The sVNT was used to analyze the binding ability of RBD to ACE2 after neutralizing RBD with antibodies in the serum. Serum collected two weeks after the final immunization was quantified according to the manufacturer’s instructions (Sugentech, CONE001E). Briefly, sera were serially diluted in dilution buffer and treated with HRP-conjugated RBD for 30 min at 37 °C. The samples were added to a plate coated with the human ACE2 protein and incubated for 15 min at 37 °C. Following incubation, the plates were washed five times with the wash solution. After the final wash, the plates were developed using TMB solution, and the reaction was arrested with a stop solution. The plates were analyzed at 450 nm using the SpectraMax Plus384 (Molecular Devices). The reciprocal of the dilution that resulted in a binding inhibition rate of 20% or more (PI20) was defined as the neutralizing antibody titer.

### IFN-γ ELISPOT

A mouse IFN-γ ELISPOT set (BD 551083) was used as directed by the manufacturer. The ELISPOT plates were coated with purified anti-mouse IFN-γ capture antibody and incubated overnight at 4 °C. Plates were washed and blocked for 2 h with RPMI + 10% FBS (R10 medium), and 5 × 10^5^ splenocytes were added to each well and stimulated for 24 h at 37 °C in 5% CO_2_ with R10 medium (negative control), concanavalin A (positive control), or specific peptide pools (2 μg/mL). Peptide pools consisted of 15-mer peptides overlapped by 11 amino acids and spanning the entire S and N proteins of SARS-CoV-2 (GenScript). After stimulation, the plates were washed and spots were developed according to the manufacturer’s instructions. The plates were scanned and counted using an AID ELISPOT reader. Spot-forming units (SFU) per million cells were calculated by subtracting the number of negative control wells.

### Statistical analysis

Data analyses were performed using GraphPad Prism 7 (GraphPad Software). Comparisons between groups were performed using a two-tailed Student’s *t*-tests. Statistical significance was set at P < 0.05.

